# Impaired functional brain-heart interplay sustains emotion dysregulation in depressed individuals

**DOI:** 10.1101/2024.03.14.585023

**Authors:** Vincenzo Catrambone, Francesca Mura, Elisabetta Patron, Claudio Gentili, Gaetano Valenza

## Abstract

Depression is a leading worldwide cause of mental disorders and disability, strongly affecting emotional processing and regulation. Its dysfunctional psycho-physiological dynamics may be part of the a nervous-system-wise symptomatology, impacting not only patients’ psyche but also significantly influencing functional cardiovascular dynamics. Therefore, depression serves as an exemplary pathological manifestation of the dysfunctional interaction between the central and autonomic nervous systems. While recent literature has been developing specific techniques to quantify such interactions, often referred to as functional Brain-Heart Interplay (BHI), the quantitative role of BHI dynamics in depression is largely unknown. This study aims to experimentally unveil BHI patterns specific to emotional regulation and processing in subjects exhibiting depressive symptoms compared to healthy controls. Results were gathered from a cohort of 72 individuals and indicate that depressive symptoms are associated with a continuous efferent central-to-peripheral hyperactivity and an afferent peripheral-to-central hypoactivity. This hypoactivity appears to be specific to negative emotional processing. This study offers novel insights into the systemic investigation of the neuro-physiological bases of depression.

## Introduction

Depression is a severe mental disorder, the symptomatology of which is described in the Diagnostic and Statistical Manual of Mental Disorders (DSM) ***American Psychiatric Association et al. (2013***). It has a general prevalence of 4.4% ***Depression (2017***), with an estimated lifetime prevalence reaching 17% ***Blazer et al. (1994)***. It represents the leading worldwide cause of disability regarding mental health and has been recognized as a risk factor for early mortality, cardiovascular disease, and neurological disease ***Van der Kooy et al. (2007***); ***Frasure-Smith and Lespérance (2005***). Depression has a significant impact on public health, being among the foremost leading causes of the global burden of disease ***Press and Geneva (2008***).

Depression, characterized by prolonged sadness and anhedonia, significantly affects people’s everyday life and well-being. It often manifests with somatic symptoms such as fatigue, loss of energy, difficulty sleeping, as well as cognitive-affective symptoms such as feelings of melancholy, guilt, pessimism, and difficulty concentrating ***Storch et al. (2004***).

The heterogeneity of somatic, cognitive, and affective symptom presentations in different forms of depression complicates diagnosis and prognosis, often leading to disagreement among clinicians ***Gentili (2017***), while objective measures to support diagnosis are not available. Therefore, characterizing the psycho-physiological features of the dysfunctions emerging in individuals with depressive symptoms may aid in overcoming this difficulty ***Cuthbert and Insel (2013***); ***Sanislow et al. (2010***). In this context, the investigation of psychophysiological reactivity to emotional stimuli to characterize the resulting altered emotional experiences has emerged as a particularly effective research topic ***Joormann and Gotlib (2010***); ***Rottenberg (2005***); ***McLaughlin et al. (2011***).

Within this paradigm, several hypotheses and methods have been proposed in an effort to shed light on the role of impaired emotional response in the early manifestation of depression symptoms ***Hill et al. (2019***); ***Kring and Bachorowski (1999***). Consistently, individuals affected by depression tend to be more sensitive to unpleasant stimuli and to focus on them for extended periods of time ***Goeleven et al. (2006***), which contributes to the maintenance of depression ***Scher et al. (2005***); ***Ingram (1984***). It should be noted that neuro-physiological and psycho-physiological investigations have shown that individuals with depressive symptoms exhibit altered responses to stimuli with both positive and negative valence ***Werner-Seidler et al. (2013***); ***Garcia et al. (2016***).

In the context of characterizing the neurophysiological dysfunctions, a plethora of research has focused on the neural dynamics evident in clinical and subclinical depression under various experimental conditions ***de Aguiar Neto and Rosa (2019***), particularly leveraging electroencephalogram (EEG)-derived features such as spectral activity ***Grin-Yatsenko et al. (2009***), connectivity ***Miljevic et al. (2023***), or symmetry metrics ***Thibodeau et al. (2006***). Nevertheless, depression is recognized as a systemic disorder extending beyond a purely neurological dimension. The Vascular Depression theory has been formulated to this extent ***Alexopoulos et al. (1997***); ***Taylor et al. (2013***), acknowledging depression as a major cause of cardiovascular disorders and as associated with a higher risk and worse prognosis of coronary heart disease ***Barth et al. (2004***). Dysfunctional cardiovascular dynamics are known to have a significant impact on the risk of depression through both direct physical effects and indirect biochemical, physiological, and behavioral pathways ***Rovai et al. (2015***). Notably, one of the most common physical comorbidities of depression is cardiovascular disease ***Glassman (2007***), and this comorbidity may initiate a downward spiral effect in which depressive and cardiac symptoms reinforce each other ***Penninx (2017***); ***Kidwell and Ellenbroek (2018***). Accordingly, several functional connections between autonomic nervous system (ANS) dynamics and depression symptoms have been identified ***Carney et al. (2005***); ***Sgoifo et al. (2015***). Specifically, individuals with major depression exhibit sympathetic hyper-tonia ***Licht et al. (2015***) and decreased vagal tone compared to healthy controls ***Kemp et al. (2010***). Vagal hypoactivity has been associated with specific symptoms of depression, including harsh behavioral responses, decreased social engagement, and somatomotor deficiencies ***Porges (2001***). Furthermore, nonlinear signal processing techniques have been able to distinguish individuals with mood disorders from healthy controls ***Licht et al. (2015***), for example, by demonstrating a significant increase in heart rate variability (HRV) complexity in the presence of subclinical depression ***Valenza et al. (2014***). However, singular HRV measures quantifying ANS activity are known to be non-specific, meaning that similar changes in HRV dynamics might be encountered and found to be correlated with different pathological and physiological conditions ***Saul and Valenza (2021***).

More generally, broader clinical data suggests that a patient’s mood, as well as brain and car-diovascular dynamics are casually related, and so mood disorders may be linked to changes in the CNS-ANS interplay, often referred to as functional Brain–Heart Interplay (BHI) ***Catrambone and Valenza (2021***); ***Taggart et al. (2011***); ***Valenza (2023***); ***Catrambone et al. (2021a***); ***Malandrone et al. (2024***); ***Verdonk et al. (2024***); ***Blickle et al. (2024***); ***Schulz et al. (2022***); ***Terhaar et al. (2012***). BHI has conceptually been linked to the Brain-Heart Axis, a functional system comprising all neural, mechanical, hormonal, and electrical communication pathways between the CNS and ANS. Recent findings highlight that BHI dynamics are highly directional ***Candia-Rivera et al. (2022***); ***Catrambone and Valenza (2021***); ***Pereira et al. (2013***); ***Catrambone et al. (2021a***); ***Malandrone et al. (2024***). In healthy individuals, on the one hand, directional BHI is modulated by stress ***Catrambone et al. (2024***); ***Candia-Rivera et al. (2023***), as well as emotional processing ***Candia-Rivera et al. (2022***), with the ascending heart-to-brain changes occurring first in time. On the other hand, directed brain-to-heart dynamics have a specific response timing with a s period in response to sympathovagal elicitation ***Catrambone et al. (2021c***). In patients affected by mood disorders, a functional hyperdrive of CNS control over the ANS has been linked to a higher risk of cardiovascular illnesses ***Pereira et al. (2013***). Accordingly, such a hyperdrive in the directional brain-to-heart interplay has been found in individuals with mild depressive symptoms at rest compared to healthy subjects ***Catrambone et al. (2021a***), whereas no differences have been detected in the opposite ascending heart-to-brain direction, particularly at the cortical level. Consistently, analyses on the heartbeat-evoked potential, a measure considered to be a marker of the central response to bodily stimuli, did not find differences between depressive subjects and healthy controls at rest ***Catrambone et al. (2021a***); ***Schulz et al. (2022***).

While the aforementioned evidence clearly suggests that functional BHI dynamics are involved in depression pathophysiology, the quantitative role of directional BHI sustaining emotion dysregulation in depression remains unclear. To this end, in this study, we investigate nervous-system-wise psychophysiological responses to emotional stimuli in subjects showing depressive symptoms. Functional BHI is quantified through physiologically-plausible modeling providing directional estimates in the time and frequency domains. Further details follow below.

## Materials and Methods

### Dataset description

All participants were recruited at the University of Padua (Padua, Italy). The experimental protocol was defined in agreement with the Declaration of Helsinki and approved by the local ethical committee (protocol no. 3727). The experimental procedure involved 72 volunteering participants (17 males, aged 22.5 ± 2.4 years). Exclusion criteria included the presence of any of the following conditions: psychotic symptoms, psychotropic medication, substance abuse, neurological disorders, and a history of brain injury. Participants were instructed to abstain from consuming alcohol the day before the experimental session and to refrain from using stimulants (such as caffeine and nicotine) on the day of the experiment. An ad-hoc interview was administered to assess sociodemographic variables (i.e., age, sex, education) and medical status (i.e., medical conditions and pharmacological therapy). Moreover, participants completed the Beck Depression Inventory, 2nd edition ***Beck et al. (1996***); ***Sica et al. (2007***) to assess the presence of depressive symptoms.

The experimental session took place in a dimly lit, soundproof room, where participants were seated on a comfortable chair while an EEG cap was applied to their scalp. Prior to the emotional passive viewing task, participants completed six practice trials, consisting of two images from each of the three emotional categories (pleasant, neutral, and unpleasant images). Following the practice trials, participants completed the passive viewing task, which consisted of a total of 72 images. Specifically, the 72 digital color photographs were selected from the International Affective Picture System (IAPS, ***Lang and Bradley (2007***)) and equally divided into three categories: pleasant (e.g., laughing faces or baby playing scenes), neutral (e.g., domestic objects), and unpleasant (e.g., animal assaults). According to the well-known circumplex model of affect ***Russell (1980***), the three considered categories could be differentiated in terms of valence (i.e., pleasantness of a stimulus), whereas pleasant and unpleasant images were similar in terms of high arousal (i.e., strength of the emotional perception), unlike the neutral pictures, which were low arousal stimuli. Operationally, a white fixation cross was presented at the center of the screen for 3 seconds preceding each image appearance, which lasted on the screen for the following 6 seconds. After each image presentation, a white fixation cross appeared again in the center of the screen for a period ranging from 6 to 8 seconds. The passive viewing task pipeline is graphically represented in figure 1.

**Figure 1.**
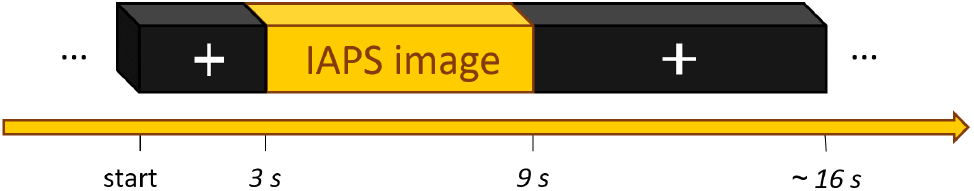
Graphical representation of the emotional picture passive viewing procedure.

Pictures were presented in a semi-randomized order, ensuring that stimuli belonging to the same emotional category were not shown consecutively. All participants signed an informed consent form and received €15 compensation for participating in the experiment.

During the experimental procedure, physiological recordings were acquired using an eego amplifier (ANT Neuro, Enschede, Netherlands) and eego™software on a computer platform. EEG signals were acquired through a flexible cap featuring 32 electrodes positioned according to the 10–20 System, with online referencing to CPz. Channel impedance was constantly maintained below 10*k*Ω, and a sampling frequency of 1000*Hz* was applied.

Additionally, the ECG was acquired with disposable Ag/AgCl electrodes positioned according to the lead II Einthoven’s configuration, and the signal was recorded with a sampling rate of 1000*Hz*. Two further electro-oculographic electrodes (EOGs) were placed to monitor eye movements and blinks.

### EEG processing

The EEG recordings were pre-processed using a semi-automated pipeline running on MATLAB R2023b (The MathWorks Inc., 2023) software and the EEGLab toolbox ***Delorme and Makeig (2004***). Briefly, the pipeline comprised standard methods, such as down-sampling to 500*Hz*, initial rereferencing to mastoids, and band-pass filtering within the range of [1, 30]*Hz*. Subsequently, bad channels were identified based on the following conditions: flatline for more than 5*s*; low correlation with nearby channels; line noise at 50*Hz* exceeding 4 standard deviations of the total signal. Following this, artifacts were detected and rejected using the artifact subspace reconstruction ***Mullen et al. (2015***) and IClabel ***Pion-Tonachini et al. (2019***) methods. Eventually, predetermined bad channels were interpolated using the spherical algorithm to match the number and location of electrodes, and an average re-referencing was performed, as suggested for BHI applications ***Candia-Rivera et al. (2021***).

Six participants’ recordings were excluded due to the presence of excessive and irreparable artifacts, leaving a total sample of 66 individual recordings.

After preprocessing, the EEG power spectral density (PSD) was extracted by applying the short-time Fourier transform with a Hamming taper. A 2*s* sliding time-window with a 0.25*s* step was employed, resulting in a spectrogram sampled at 4*Hz* in the time domain and at 0.5*Hz* in the frequency domain. The four canonical frequency bands (delta: δ ∈ [1 − 4]*Hz*, theta: θ ∈ [4 − 8]*Hz*, alpha: α ∈ [8 − 12]*Hz*, and beta: β ∈ [12 − 30]*Hz*) were then obtained by integrating the PSD series within the respective frequency ranges.

### Heart Rate Variability processing

HRV series were obtained by applying a R-peak detection algorithm on the ECG series, using the well-known Pan-Tompkins method. The ECG signals were initially bandpass filtered in the range of [0.5, 25]*Hz*. The obtained series, along with the interbeat interval histogram, underwent visual inspection performed by experts. Corrections to remove physiological and algorithmic artifacts were made where necessary through the Kubios software ***Tarvainen et al. (2014***).

The smoothed pseudo-Wigner-Ville distribution method was employed to extract the time-resolved spectrogram of the HRV series, which was then integrated into the canonical HRV band ranges: the low-frequency (LF) band in the interval of [0.04, 0.15]*Hz*, and the high-frequency (HF) band in the range of [0.15, 0.4]*Hz*. Specifically, although some controversies exist ***Valenza et al. (2018***), this spectral representation is widely used in the literature: the LF band is considered a non-specific marker of sympathovagal activity, while the HF band represents parasympathetic activity ***Rajendra Acharya et al. (2006***). The two time-varying spectral estimations derived from HRV (i.e., LF and HF) were then sampled at 4*Hz* to maintain consistency with those derived from the EEG processing.

### Brain–heart interplay estimation

BHI measures were derived using the synthetic data generation (SDG) model presented in ***Catrambone et al. (2019***, 2021b). Briefly, the SDG model is based on the computational representation of two coupled models which separately represent the EEG- and HRV-derived series. In particular, the EEG recordings are modeled as a multi-oscillator system (i.e., there is one oscillator for each frequency band considered) whose amplitudes are singularly described through an exogenous autoregressive (XAR) process of the first order ***Al-Nashash et al. (2004***). Each XAR process describes, through its exogenous term, the time-resolved ascending interplay going from the heart to the brain, specific for that specific oscillation.

Complementarily, the computational model describing the cardiovascular dynamics extends the widely used integral pulse frequency modulation (IPFM) model proposed by ***Brennan et al. (2002***), which conceives the HRV series as given by an integrator firing when the sympathovagal activity reaches a determined threshold. Here, the IPFM model accounts for the interplay going from the brain to the heart through the introduction of an exogenous term driving sympathovagal activity.

More generally, the fundamental concept behind the SDG model is that the activity of the two underlying systems (i.e., CNS and ANS), represented by the associated electro-physiological signals (i.e., EEG and HRV, respectively) are not independent of one another, and the coupling terms try to capture these relationships in formal terms. To summarize, for combinations of EEG- and HRV-frequency components, time-varying directional BHI biomarkers represent an instantaneous assessment of heart-to-brain and brain-to-heart interactions, separately. Thus, formally, if there is an increase in a given *F*-band of the HRV spectrum at time *t*_*n*−1_, and a proportional increase in a certain *B*-band of the EEG spectrum at time *t*_*n*_, then the SDG model would capture a positive value of *C*_*F →B*_(*t*_*n*_). In other words, the activity of a specific frequency range of the HRV series is positively linked to the activity of a specific frequency range of the EEG series, thus representing a functional relationship between the two electro-physiological recordings, which in turn is a representation of a functional relationship between the two physiological systems.

The quantitative estimation of the family of BHI biomarkers derived through the SDG model is detailed in ***Catrambone et al. (2019***), and a freely available open source MATLAB implementation can be found at^1^. Eventually, the functional directional BHI indices were derived employing the SDG framework, and specifically involved separate analysis for the EEG-PSD series in the *δ, θ, α*, and *β* bands, as well as for the HRV-PSD series in the *LF* and *HF* bands. We remark that the SDG model allows estimating functional BHI in a time-resolved structure that follows the time resolution of the PSDs given as input to the model, thus allowing the obtained BHI indexes to be sampled at 4*Hz*. All the derived BHI indices are reported in table 1.

**Table 1.** Brain-Heart Interplay indices.

| BHI | | $\delta$ | $\theta$ | $\alpha$ | $\beta$ |
| --- | --- | --- | --- | --- | --- |
| brain-to-heart | LF | $C_{\delta \rightarrow LF}$ | $C_{\theta \rightarrow LF}$ | $C_{\alpha \rightarrow LF}$ | $C_{\beta \rightarrow LF}$ |
| | HF | $C_{\delta \rightarrow HF}$ | $C_{\theta \rightarrow HF}$ | $C_{\alpha \rightarrow HF}$ | $C_{\beta \rightarrow HF}$ |
| heart-to-brain | LF | $C_{LF \rightarrow \delta}$ | $C_{LF \rightarrow \theta}$ | $C_{LF \rightarrow \alpha}$ | $C_{LF \rightarrow \beta}$ |
| | HF | $C_{HF \rightarrow \delta}$ | $C_{HF \rightarrow \theta}$ | $C_{HF \rightarrow \alpha}$ | $C_{HF \rightarrow \beta}$ |

### Statistical analysis

Group-wise statistical analysis on arousal and valence levels self-reported by participants was performed through the non-parametric one-way Friedman test to statistically assess whether the designed experimental procedure effectively elicited the intended emotional responses.

To assess subject-wise functional BHI indexes for the different emotional experimental conditions, a BHI time average was performed for each visual stimulus presented to each subject. Subsequently, a further averaging was conducted among all the BHI indexes elicited by visual stimuli that were homogeneous in terms of arousal and valence. For instance, all BHIs measured during visual emotional stimuli with positive valence and high arousal (POS) were averaged. This procedure resulted in a subject-specific single functional BHI measure for each channel and combination of EEG- and HRV-derived frequency bands, separately for POS, neutral valence and low-arousal (NEUT), and high arousal and negative valence (NEG) visual emotional elicitation.

The aforementioned measures were first employed to statistically test the null hypothesis of no BHI differences between two experimental cohorts: a healthy controls (HC) group and a group of subjects with depressive symptoms (DS), across different emotional stimuli, using the non-parametric Mann-Whitney test for independent samples. Second, we statistically tested the null hypothesis of no differences among the emotional stimuli in the group with depressive symptoms, using the non-parametric Wilcoxon test for paired samples. A significance threshold of *α* = 0.05 was considered. This statistical analysis was performed separately for the combinations of BHI directions, EEG- and HRV-derived frequency bands, for all the 30 EEG channels. This procedure raised the issue of multiple comparisons, which we addressed by employing a spatial correction through permutation procedure ***Friston et al. (1994***), assessing a minimum cluster size of four electrodes being concurrently significant to reject the null hypothesis.

## Experimental Results

### The experimental protocol induced an emotional elicitation

Experimental results on the self-reported arousal scores revealed significant differences (p-value < 0.001), and a further post-hoc test verified significantly higher perceived arousal for both pleasant and unpleasant stimuli compared to neutral ones (both with p-value < 0.001). No differences were identified between arousal levels elicited by pleasant and unpleasant pictures (p-value = 0.58). Similarly, the statistical analysis performed on the self-reported valence level reported a statistically significant group-wise effect (p-value < 0.001). This was further verified by a post-hoc test, which highlighted more negative levels of valence for unpleasant stimuli compared to both neutral (p-value < 0.001) and pleasant ones (p-value < 0.001), as well as more positive valence scores for pleasant images compared to the neutral pictures (p-value < 0.001).

### Presence of depressive symptoms

The average BDI-II score was 13.6 with a standard deviation of 10.5. Among the 66 subjects, 34 reported absent to minimal depressive symptoms (BDI-II score in the interval [0 - 11]), 19 reported mild depressive symptoms (BDI-II score in the range [12 - 19]), 8 showed moderate depressive symptoms (BDI-II in the interval [20 - 28]), and 5 had severe depression symptoms (BDI-II in the range [29 - 63]). To differentiate between healthy controls and subjects with depressive symptoms, we divided the cohort into two groups based on the BDI-II score. We chose a BDI-II index equal to 12 as the threshold, meaning that subjects with BDI-II scores in the interval [0-11] were assigned to the healthy control (HC) group, and participants reporting a BDI-II score equal to or higher than 12 were assigned to the group of subjects with depressive symptoms (DS), following the Italian validation of the BDI-II.

For completeness, we report the average BDI somatic score, which was 8.36 with a standard deviation of 6.01 (i.e., BDI-II-S score ranges from 0 to 36), and the average BDI cognitive-affective index, which was 5.22 with a standard deviation of 5.15 (i.e., BDI-II-C scores range from 0 to 27). The presence of depressive symptoms did not significantly affect the emotional elicitation induced by the experimental protocol. Results are presented in Table 2, where average arousal and valence levels are reported for each experimental group across all three stimulus classes (i.e., positive, neutral, and negative images). A statistical analysis performed through the non-parametric Mann-Whitney test for independent samples comparing arousal and valence levels in the two experimental cohorts found that, although a tendency exists in terms of divergent perceived valence in negative images (p-value equals to 0.081), no significant differences were observed in perceived arousal or valence.

**Table 2.**
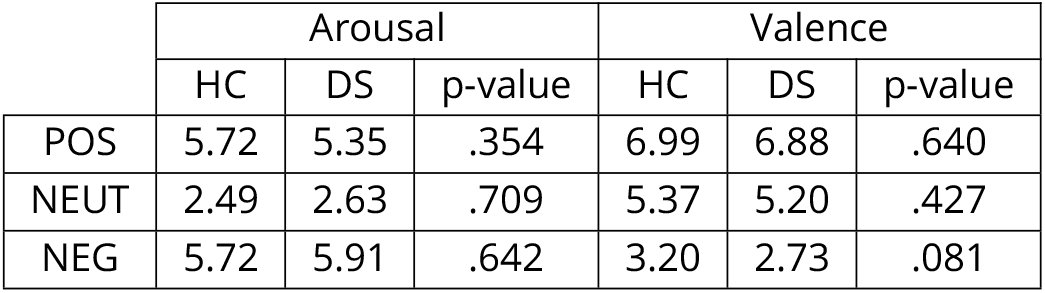

### Inter-group HC vs. DS differences in functional BHI

Experimental results related to BHI estimates are presented as topographical representations of statistically significant electrodes, utilizing a color-code where the intensity of color is inversely proportional to the logarithm of the achieved p-values in the statistical comparison. Essentially, the higher the color intensity, the lower the magnitude of the sub-threshold corrected p-value.

Specifically, in Figure 2, statistical results obtained through the application of the Mann-Whitney non-parametric test for independent samples are depicted. This compares the average BHI elicited by neutral images in HC and DS groups. The color-code provides information about the sign of the statistics. The blue region highlighted in the *C*_*θ*→*HF*_ case represents a large fronto-temporal centro-right cluster of significant differences between the two experimental groups, where the DS group exhibits a higher BHI measure compared to the HC group. No other significant differences are reported when averaging the emotional picture with neutral (i.e., low arousal and negligible valence) in terms of functional BHI in any of the combinations of EEG- and HRV-derived frequency bands.

**Figure 2.**
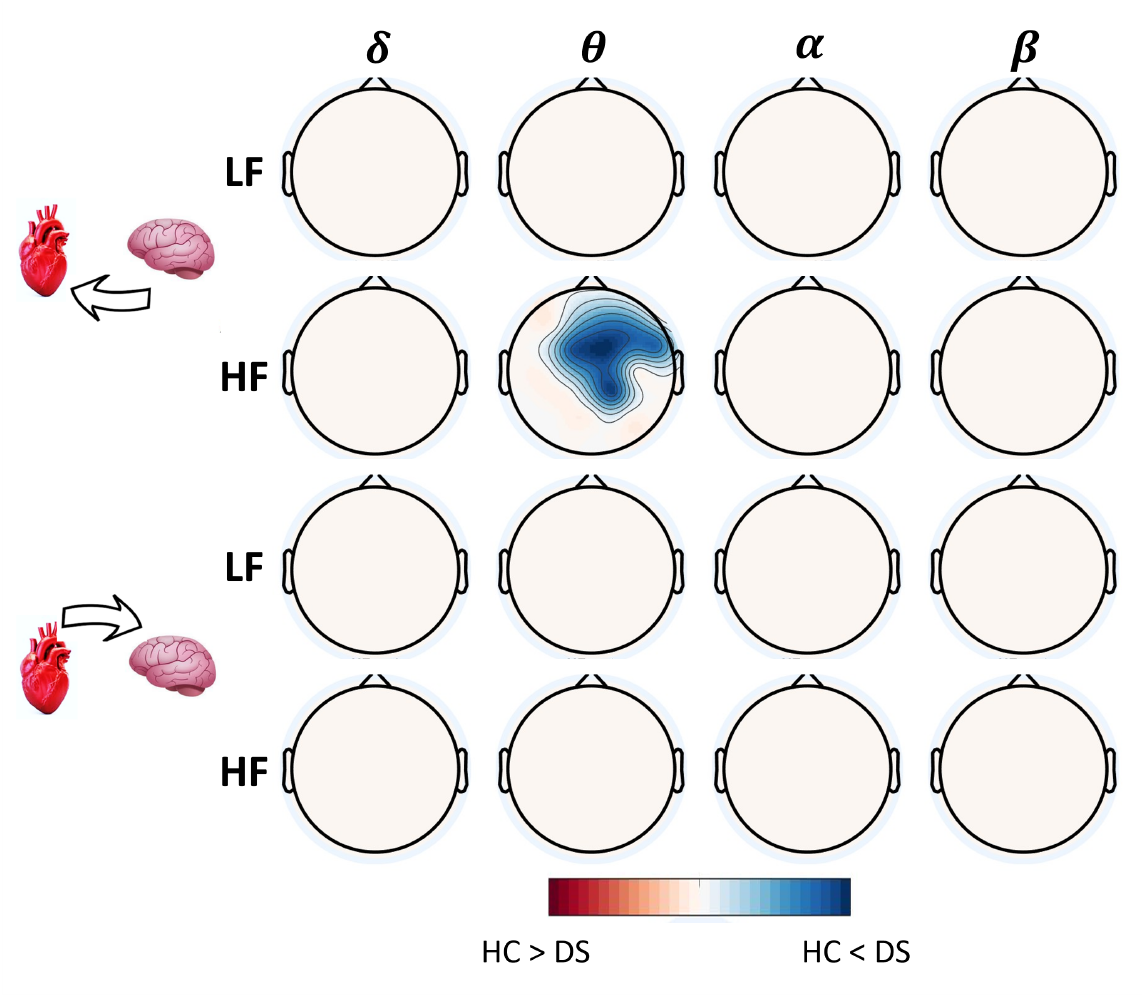
Topographical representation of statistical results obtained by the application of the Mann-Whitney non-parametric test for independent samples, comparing the average BHI elicited by neutral images in healthy controls (HC) and subjects with depressive symptoms (DS). The first two rows refer to the directional brain-to-heart interplay in HRV-LF (first row) and HF (second row), respectively. The last two rows refer to the directional heart-to-brain interplay in HRV-LF (third row) and HF (fourth row), respectively. The four columns refer to the EEG-derived frequency bands (i.e., *δ, θ, α*, and *β*). White regions represent non-significant comparisons, whereas areas are colored when significant differences were detected in the corresponding electrodes. These regions are shown in red when the HC sample median was greater than DS’, and blue in the opposite case.

In Figure 3, statistical results obtained from the application of the Mann-Whitney non-parametric test for independent samples are presented, comparing the average BHI elicited by negative images (left subpanel (a)) and positive images (right subpanel (b)) in HC and DS groups. Remarkably, significant results are observed with NEG stimuli, denoting images with high arousal and negative valence, where a notable brain-to-heart component is evident. Here, the BHI estimates are significantly higher in the DS group compared to the HC group. This is particularly emphasized in a bounded right fronto-temporal region considering *C*_*θ*→*LF*_, whereas for *C*_*α*→*LF*_, a broad area comprising the central region of the scalp, as well as centro-posterior, left temporal, and right frontal areas, is notable.

**Figure 3.**
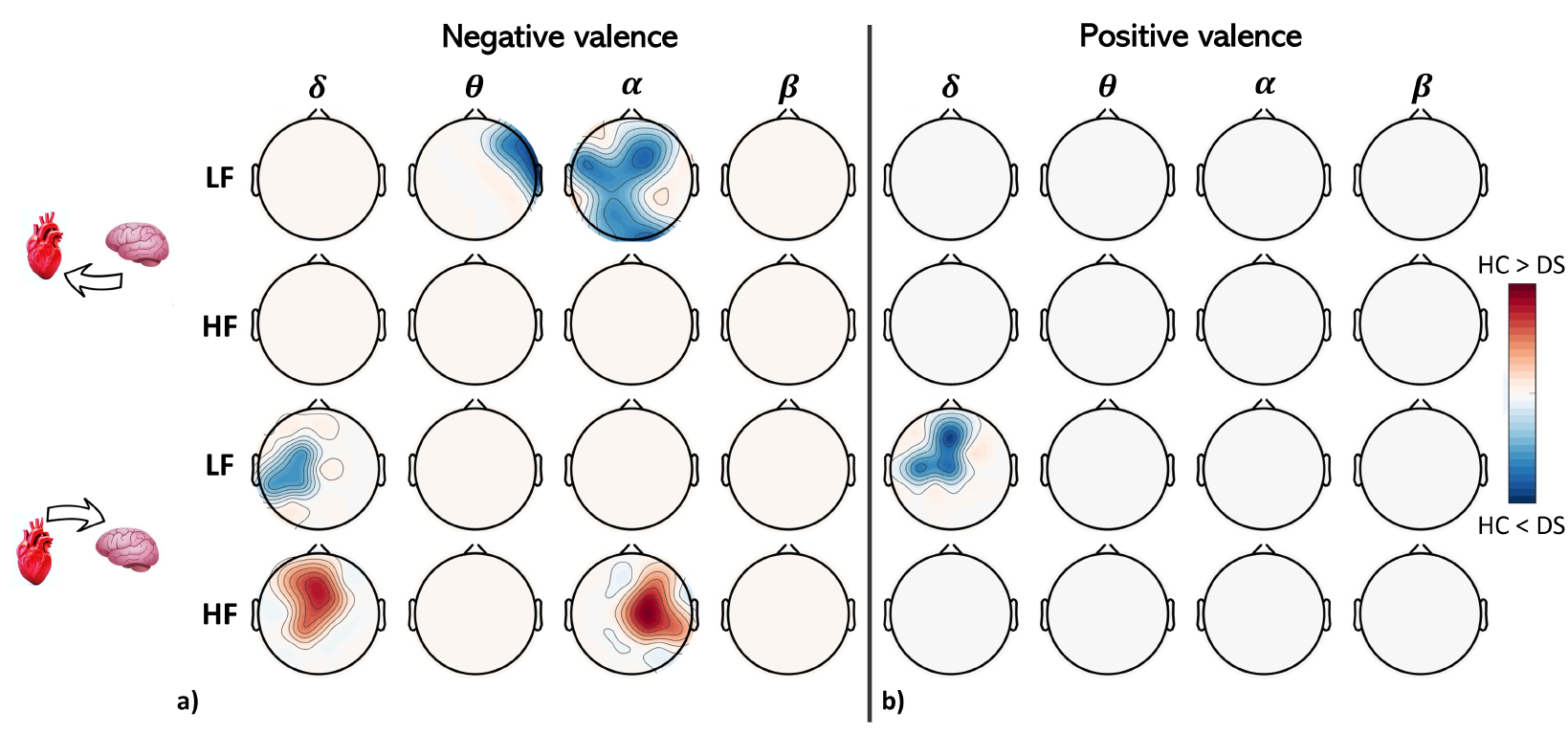
Topographical representation of statistical results obtained by the application of the Mann-Whitney non-parametric test for independent samples, comparing the average BHI elicited by negative images (left subpanel (a)) and positive images (right subpanel (b)) in healthy controls (HC) and subjects with depressive symptoms (DS). The first two rows refer to the directional brain-to-heart interplay in HRV-LF (first row) and HF (second row), respectively. The last two rows refer to the directional heart-to-brain interplay in HRV-LF (third row) and HF (fourth row), respectively. The four columns refer to the EEG-derived frequency bands (i.e., *δ, θ, α*, and *β*). White regions represent non-significant comparisons, whereas areas are colored when significant differences were detected in the corresponding electrodes. These regions are shown in red when the HC sample was greater than DS’, and blue in the opposite case.

Conversely, in the heart-to-brain component, the comparison yields the opposite sign, indicating that the BHI is higher on average in the HC group compared to the DS group. This is especially evident for the HRV-HF band on one side and the EEG-δ and α bands on the other. Specifically, the comparison on *C*_*HF* →*δ*_ highlights a centro-frontal significant area, while on *C*_*HF* →*α*_, the significant region is more lateralized on the right centro-temporal hemisphere. Moreover, a central region, slightly lateralized on the left side, is found to be statistically significant in the comparison involving *C*_*LF*→*δ*_, with BHI estimates being higher in the DS group compared to the HC group, for both negative (Figure 3.a) and positive (Figure 3.b) stimuli. No further significant regions have been detected.

### Intra-group differences in functional BHI

Figure 4.a shows statistical results obtained by the application of the Friedman non-parametric test for paired samples, comparing the average BHI elicited by images with three different emotional contents (i.e., POS, NEG, and NEUT) in the DS group. The significant results are direction-specific in the context of BHI, meaning that all the highlighted interactions belong to the descending brain-to-heart interplay, particularly involving the sympathovagal HRV-LF band and the entire EEG spectrum, even with important differences in terms of scalp regions. More in detail, the *C*_*δ*→*LF*_ interplay has been found to be significantly different in the three emotional picture stimuli, on average, in a centro-frontal area occupying the inter-hemisphere midline and partially inclined on the right hemisphere. The *C*_*θ*→*LF*_ interplay drawn in the first row and second column of Figure 4.a reports significant differences in the DS group located in the right hemisphere in the anterior and central regions. Further significant differences have been detected analyzing the *C*_*α*→*LF*_ interplay, particularly in a central horizontal stripe involving left temporal, central, and right temporal regions, with higher significance reached in the left centro-temporal area. Eventually, a small centro-occipital scalp area is highlighted to have significant different *C*_*β*→*LF*_ in the top-right corner of Figure 4.a. No significant differences have been detected in the descending brain-to-HF, as well as in the ascending heart-to-brain interplay, independently from the frequency band considered. Conversely, the analogous statistical analysis performed on BHI measures gathered from the HC group reported in Figure 4.b shows that in healthy subjects, the BHI changes elicited by the group-wise analysis are significant in the opposite ascending heart-to-brain combination, specifically in the *C*_*HF →*δ_ case in a central scalp region.

**Figure 4.**
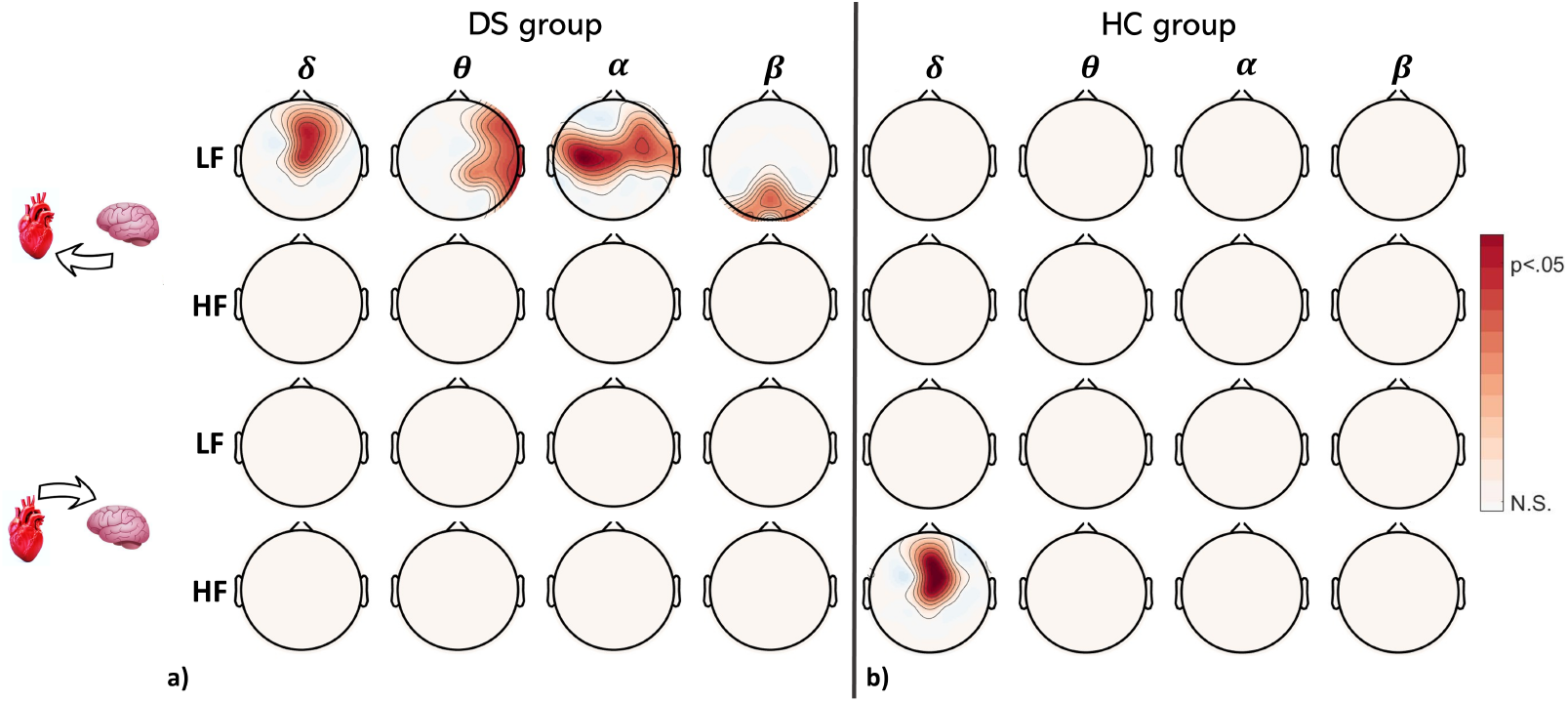
Topographical representation of statistical results obtained by the application of the Friedman non-parametric test for paired samples, comparing the average BHI elicited by images with three different emotional contents (i.e., POS, NEG, and NEUT), in subjects with depressive symptoms (left subpanel (a)) and healthy controls (right subpanel (b)). The first two rows refer to the directional brain-to-heart interplay in HRV-LF (first row) and HF (second row), respectively. The last two rows refer to the directional heart-to-brain interplay in HRV-LF (third row) and HF (fourth row), respectively. The four columns refer to the EEG-derived frequency bands (i.e., *δ, θ, α*, and *β*). White regions represent non-significant comparisons, whereas red regions highlight significant differences.

## Discussion

We investigated the nervous-system-wise psychophysiological responses elicited by different emotional stimuli in subjects with depressive symptoms, comparing them to those measured in healthy controls. We estimated the directional functional BHI through a synthetic data generation model, quantifying ascending heart-to-brain and descending brain-to-heart interactions in specific EEG- and HRV-derived frequency ranges, separately.

The analyzed data were gathered from a final cohort comprising 66 subjects, with 34 belonging to the HC group and 32 to the DS group. Emotional elicitation was induced by presenting standard images with fixed levels of arousal and valence, i.e., neutral (low arousal, neutral valence), positive (high arousal, positive valence), and negative (high arousal, negative valence) emotional pictures. A statistical evaluation of self-assessment manikin scores confirmed that the visual image stimulation effectively elicited an emotional response in the experimental cohort. Such an emotional response did not significantly differ between the two groups, i.e., no significant differences were detected in the perceived arousal and valence between healthy controls and subjects with depressive symptoms, neither in the positive, neutral, nor negative images (see tab. 2). This is in line with the literature on subclinical (Messerotti et al., 2019; Sloan et al., 2010) and clinical depression (Dichter et al., 2004), where self-reported measures of valence and arousal were not linked with depressive symptoms. Overall, this suggests that differences in BHI are detectable even when they do not manifest in self-reported valence and arousal. As a consequence, it might be possible that functional BHI estimates reveal underlying cortical processes that conscious subjective ratings of emotional experiences do not distinguish. These findings emphasize the role of adding psychophysiological measurements to clinical and research trials in the study of patterns associated with depression.

In terms of the BHI differences detected in the two experimental groups (i.e., subjects with depressive symptoms and healthy controls), the present study highlights significant insights into the characterization of emotional processing and dysregulation regulation experienced by depressive patients. Indeed, an efferent brain-to-heart hyperdrive, particularly considering the EEG-*θ* and HRV-HF frequency bands, in the DS group was reported during neutral stimulation (see Figure 2). This result is in agreement with an enhanced descending BHI reported in subjects with mild depressive symptoms in a resting state scenario ***Catrambone et al. (2021a***). Nevertheless, while in the resting state of dysphoric subjects an over-activation of descending BHI was detected in the HRV-LF sympathovagal band, here the experimental results reported in Figure 2 confirm the general directional BHI changes but highlight the HRV-HF vagal band. This difference might be associated with the aspecificity of the spectral components in which the heartbeat dynamics spectrum has been divided (i.e., LF, HF); in fact, the vagal activity is known to significantly affect the power in the HRV LF band ***Valenza et al. (2018***). Moreover, we observed a descending BHI hyperdrive in the *C*_*θ*→*LF*_ and *C*_*α*→*LF*_ functional directions during the processing of negative emotional stimuli (i.e., images with negative valence) in subjects with depressive symptoms compared to healthy controls (see Figure 3.a). Such changes were associated with a hypoactivation of ascending BHI in *C*_*HF →*δ_ and *C*_*HF →*α_ combinations. Interestingly, in healthy subjects, the ascending HF-to-brain interplay has been found to be the fastest neuro-cardiovascular response to emotional elicitation, anticipating the following brain-to-heart response in the opposite direction ***Candia-Rivera et al. (2022***). It must be noted that most of these differences do not arise in Figure 3.b, related to the differences occurred in stimulation via images with positive valence. In this case, the only significant BHI highlighted is the *C*_*LF→*δ_ combination, located in a centro-frontal scalp region, where the DS group appears to have higher ascending interaction. The same significant BHI changes, comprising the same frequency bands in the ascending direction, were also detected during negative-valence stimuli in Figure 3.a, in a more lateralized left central region. This result may lead to the conjecture that stimuli with an emotional content of negative valence may have a higher effect on the neuro-physiological response captured by BHI estimations in subjects with depressive symptoms.

Moreover, at a speculative level, the results reported in Figure 3, particularly those associated with the inter-group differences elicited by emotional images with negative valence, might be interpreted as the concurrent manifestation of separate effects. The first being a general higher central control over cardiovascular dynamics exerted in subjects with depressive symptoms with respect to healthy controls, previously reported in the resting state, and that remains in neutral and negative emotional elicitation. The second being an impairment of the healthy cardiac driving over the central emotional processing and regulation mainly mediated by the ascending HF-to-brain interplay. Indeed, a decreased central response to bodily stimuli, such as the heartbeat evoked potential, was already detected in depressive patients compared to healthy controls ***Terhaar et al. (2012***).

Focusing on the dynamical changes in BHI elicited by emotional images within the DS group (see Figure 4.a), this study reported a group-wise difference mainly attributable to the descending central control over sympathovagal activity. This seems in contrast to what is exhibited by the HC group, which, consistent with literature findings, mainly shows significant heart-to-brain changes in the first seconds of emotional elicitation ***Candia-Rivera et al. (2022***). These results further strengthen our understanding of impaired BHI in depressive subjects during emotional processing and perception, further reinforcing the association found between rumination, other depressive symptoms, or meditation and sympathovagal dysfunctions ***Tang et al. (2009***); ***Carnevali et al. (2018***); ***Ottaviani et al. (2016***).

This work comes with limitations. While functional BHI may comprise mechanical and hormonal pathways, we focused on the cortical BHI that quantifies neural-electrical activities through EEG and HRV recordings exclusively. Confounding factors such as respiration and blood pressure were not taken into account. Secondly, the aspecificity of the EEG and HRV spectral representations is also a limitation of this study. Moreover, while the emotional elicitation protocol via image presentation allows for a good estimation of the average BHI elicited in multiple repetitions, it lacks the possibility to elicit more enduring and complex emotions that would require longer video presentations. Eventually, it is widely known that depression is a very multifaceted disorder, and investigating general neuro-physiological correlates of depression might miss some effects related to specific depressive symptoms.

## Conclusion

In conclusion, depression is characterized by impaired directional functional BHI, which dynamically sustains the disorder in the absence of emotional stimuli, as well as being specific to emotional processing and dysregulation. Particularly, while depressive states seem sustained by CNS-to-ANS hyperdrive, specific ANS-to-CNS hypoactivity appears specifically associated with the processing of negative stimuli in depressed individuals. These patterns might represent a fingerprint of depressive symptoms, explaining also the higher cardiovascular risk in depression. Further studies would confirm whether these patterns are similar in clinically depressed patients. Finally, our study underscores how depressive symptoms do not only affect brain functioning but also express a systemic condition rather than an organ pathology.

## Footnotes

1 https://github.com/CatramboneVincenzo/Brain-Heart-Interaction-Indexes

## Notes

### Competing Interest Statement

The authors have declared no competing interest.

